# Comprehensive analysis of the microbial consortium in the culture of flagellate *Monocercomonoides exilis*

**DOI:** 10.1101/2024.02.02.578639

**Authors:** Alejandro Jiménez-González, Sebastian Cristian Treitli, Priscila Peña-Diaz, Anna Janovská, Vladimír Beneš, Petr Žáček, Vladimír Hampl

**Author notes:** Corresponding author: Vladimir Hampl.

## Abstract

*Monocercomonoides exilis* is the only known amitochondriate eukaryote, making it an excellent model for studying the implications of mitochondrial reduction from a cellular and evolutionary point of view. Although *M. exilis* is an endobiotic heterotroph, it can grow *in vitro*, albeit with an uncharacterized and complex prokaryotic community. All attempts to grow *M. exilis* axenically have been unsuccessful. Here, we use metagenomic sequencing at different time points during culture growth to describe the composition and dynamics of this community. We assembled genomes of 24 from at least the 30 different bacterial species within. Based on DNA read abundances, *M. exilis* represents less than 1.5%, and the representation of dominant bacterial members changes over time. Genome-scale metabolic reconstruction, differential expression analysis and measurements of metabolites in the media showed that the community depends on organic carbon oxidation, fermentation, and hydrogen production without methanogenesis. This is consistent with the rapid decline of amino acids, nucleotides, glyceraldehyde, lactate, fatty acids, and alcohols in the media. The community depends on recycling the external supply of amino acids since it has a limited capacity to fix nitrogen gas and lacks ammonia oxidizers to close the nitrogen cycle. With the senescence of the culture, we observe changes in the expression of several metabolic pathways in *M. exilis*, particularly those adapting to starvation. We do not reveal any clear metabolic link to explain the dependence of *M. exilis* on prokaryotes.

## Introduction

The gut microbiota refers to the community of microorganisms inhabiting the gastrointestinal system. Several studies have proved the influence of this community on human health (Fan and Pedersen, 2021; Jandhyala et al., 2015; Thursby and Juge, 2017), but these have mainly focused on the prokaryotic component since this represents the majority of cells and biomass. Yet, the animal gut microbiota also comprises viruses and eukaryotes, including unicellular protists. The effect of protists on the function and processes in the intestine, as well as their metabolic interactions with prokaryotes, are essentially unknown, except for some medically and veterinary important parasites, e.g.*, Entamoeba histolytica, Giardia intestinalis, Cryptosporidium* spp., *Spironucleus salmonicida, Blastocystis* spp. (Partida-Rodriguez et al., 2021; von Huth et al., 2021). However, these represent only a fraction of the intestinal protist diversity and their presence often leads to pathological states that significantly depart from the conditions found in the healthy gut.

Many intestinal protists can be successfully maintained in xenic cultures using a medium such as TYSGM-9 (Diamond, 1982; Hamann et al., 2016) and for many, this is the only feasible method of *in vitro* maintenance, given that the process of axenisation, i.e., removal of other unwanted organisms, has not been achieved. Xenic cultivation depends on establishing a stable community in culture, typically derived from the original sample, i.e., gut content or stool, which can be maintained for years by regular transfers. Whereas the community may be derived from the sampled environment, its eventual composition will most probably reduce as it adapts to the culture conditions (nutrient source, oxygen concentration, etc.) and thus will only distantly resemble the situation in the host gut. Still, the culture represents a simple and tractable model system from which information about these complex communities may be obtained. To our knowledge, such attempts are rare (Hamann et al., 2017).

In the present study, we aim to characterize the culture community derived from the stool of *Chinchilla lanigera*. It contains a single eukaryote, the commensal bacterivorous flagellate *Monocercomonoides exilis*, for which a quality genome draft is available and functionally annotated (Karnkowska et al., 2019, 2016; Treitli et al., 2021). This species belongs to Oxymonadida (Preaxostyla, Metamonada), known as inhabitants of animal intestines (Hampl, 2017; Treitli et al., 2018). The whole group, including *M. exilis*, has lost mitochondria, which is unique among eukaryotes and raises interest in their biochemistry and physiology (Karnkowska et al., 2016; Novák et al., 2023). While other oxymonad species harbor symbionts (Hampl, 2017) and specific roles and interactions have been partially elucidated, e.g. *Streblomastix strix*, a protist that takes part in the complex community of the hindgut of termites and is involved in cellulose digestion (Treitli et al., 2019), no symbiotic interaction is known for *M. exilis*. Light and electron microscopy has demonstrated that species of *Monocercomonoides* actively feed on prokaryotes (Treitli et al., 2018), hence acting as predators within the intestinal microbiota. The effect and scale of this grazing on the prokaryotic community and the level of prey selectivity are unknown.

We used a multi-omic approach to obtain a first insight into the processes of this multipartite culture community over five days of growth. Namely, we determined the species composition and changes in species abundances through time, predicted the metabolism pathways, followed gene expression changes of the most abundant members as the culture aged, and finally, measured the composition and changes of selected metabolites in the media. With all these data in hand, we attempted to reconstruct the major community biochemical processes and searched for potential interactions between its members.

## Results

### Growth of the culture and sampling of DNA, RNA and media

The development of the culture was followed for seven days in an experiment performed in triplicates (Figure 1). An aliquot of the bacterized medium, i.e., medium in which *Citrobacter portucalensis* grew overnight, representing day 0, and aliquots of days 2, 3, and 5 representing the exponential, stationary, and decline phases of *M. exilis* growth, were chosen to sample metagenomic, metatranscriptomic, and metabolomic data.

**Figure 1:**
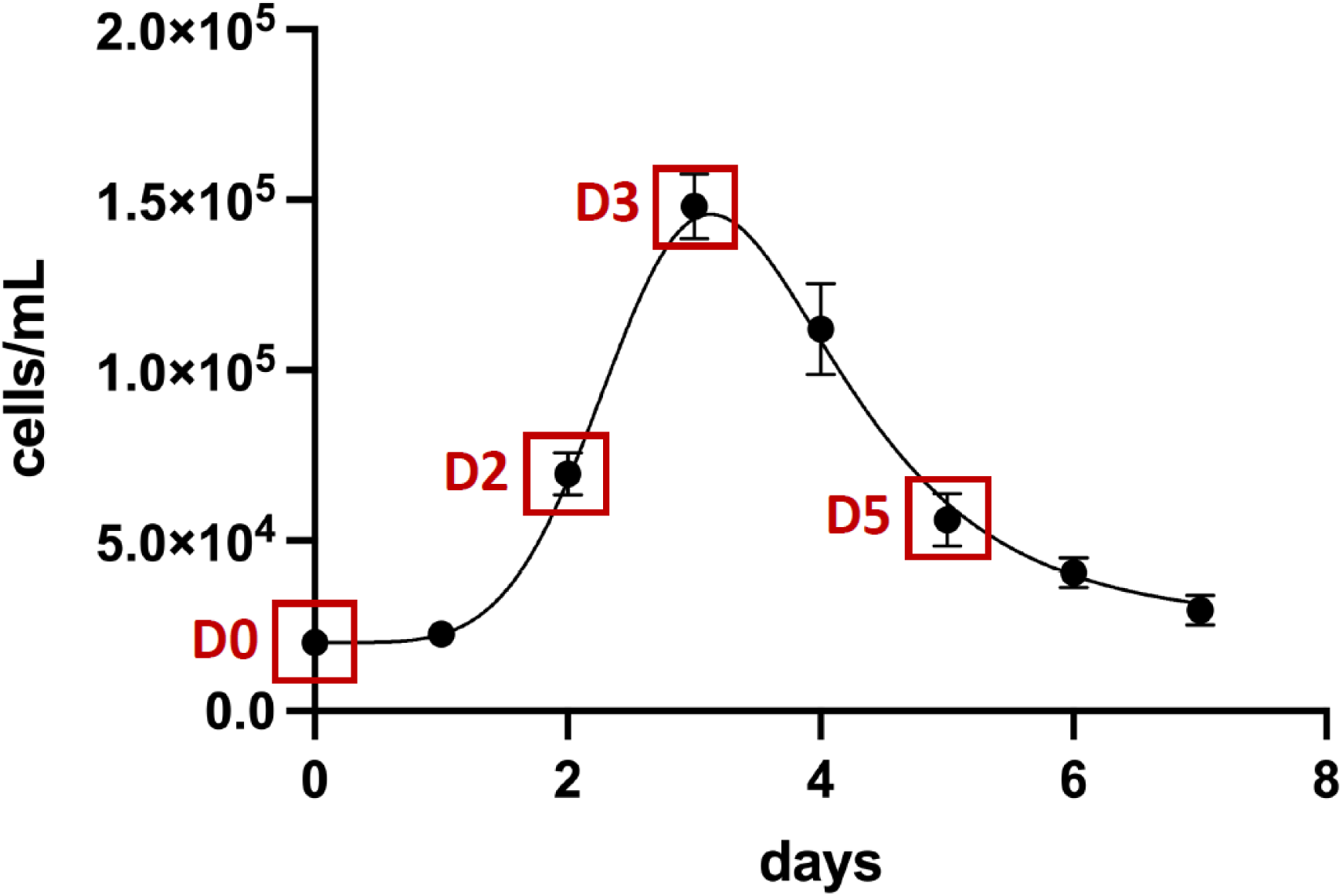
Growth curve of *Monocercomonoides exilis* and sampling points. Concentration of *M. exilis* (cells/mL^-1^) grown in TYSGM-9 medium for seven days. Samples for thorough analysis were taken on days 0, 2, 3 and 5, as highlighted by red squares. Error bars provide standard deviations of the values from three measurements.

### Community composition

Using metagenomic reads generated from triplicates of each sampling point, we assembled 24 metagenome-assembled genomes (MAGs) corresponding to the prokaryotic members of the culture community (Figure 2, Supplementary Table 1). In addition to the assembled MAGs, we identified six additional bacterial species based on the presence of 16S rRNA genes. However, their read abundances were too low to assemble them into clean MAGs (Supplementary Table 1). No viral or plasmid DNA was identified in the metagenomic data. This suggests that the community consists of one eukaryote and at least 30 bacterial species, representing seven phyla, without any archaeal or viral component.

**Figure 2:**
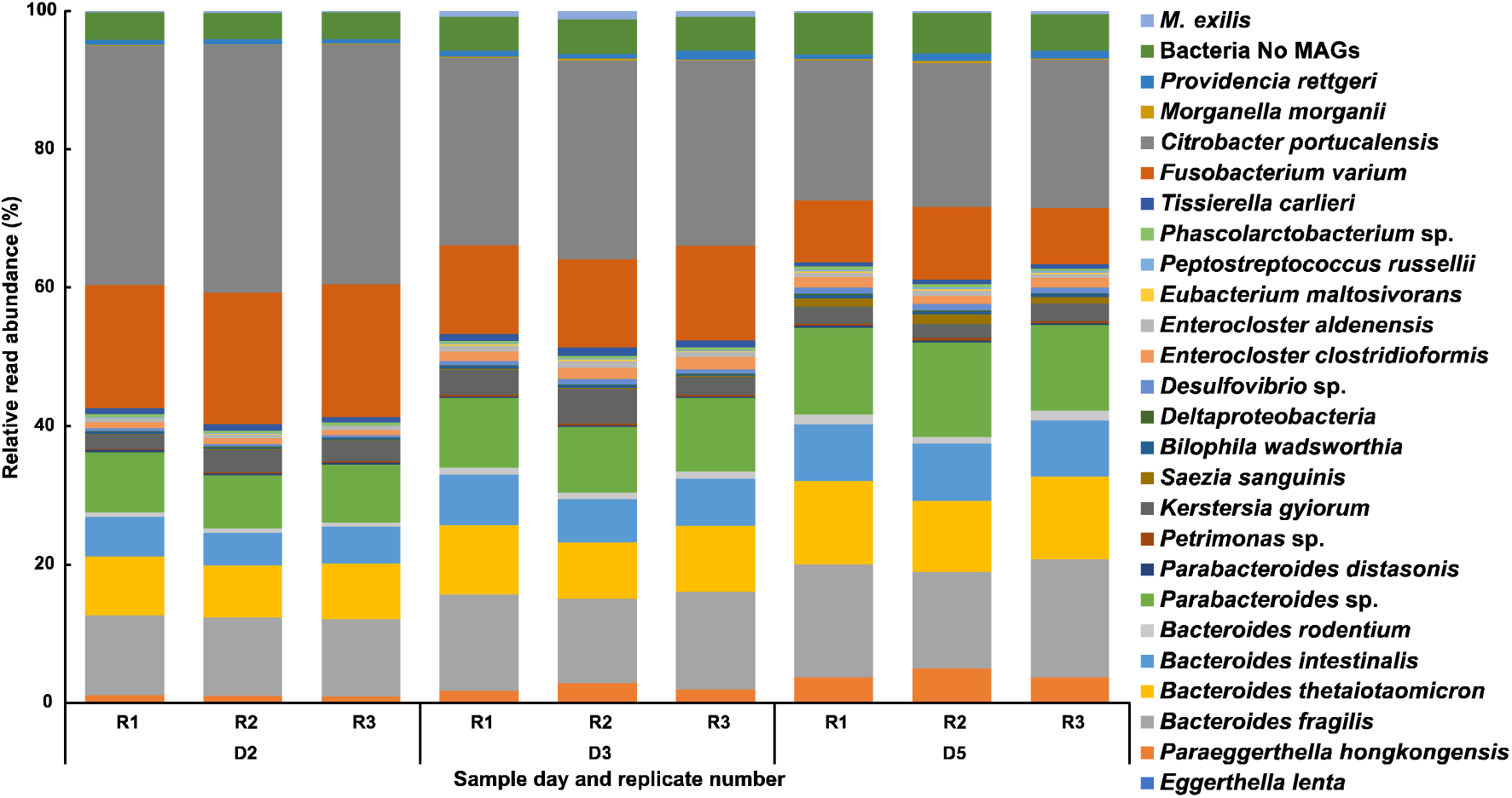
Relative read abundance of the members of the community. Each member of the community is represented as relative read abundance (%) corresponding to its genome on each day and replicate.

The quality of the 24 assembled MAGs varied, with the lowest completeness being around 83% and the vast majority having genome completeness greater than 95% (Supplementary Table 1). The estimated contamination levels are overall lower than 5%, except for *Deltaproteobacteria* sp. MAG, whose contamination was estimated to be 11.8% (Supplementary Table 1). This MAG was the least abundant during the whole experiment. We calculated the relative read abundance of each bacterium and *M. exilis* for days 2, 3, and 5 of culture. While *M. exilis* represented only 0.25% to 1.23% of the reads, the seven most abundant bacterial species comprised 80 – 90% of the reads, and the remaining bacterial species never exceeded 1% of the reads per species (Figure 2 and Supplementary Table 1). We observed a switch in the dominant groups throughout the culture growth. Not considering *Citrobacter portucalensis* (Gammaproteobacteria), which was present in the medium before inoculation of the community (“bacterization” of the medium), the most abundant bacterium on day two was *Fusobacterium varium* (Fusobacteriota). On days three and five, the abundance of this bacterium decreased, and representatives of the phylum Bacteroidota dominated. *M. exilis* reached its peak of 1.23% on day 3 (Figure 2).

### Monocercomonoides exilis metabolic capacities

Our analysis of *M. exilis* metabolism identified 933 enzymatic reactions, 31 transport and 148 enzymatic pathways (Supplementary Figure 1). As previously published (Karnkowska et al., 2019, 2016; Treitli et al., 2021), *M. exilis* is equipped with a limited repertoire for energy metabolism relying on glycolysis to produce pyruvate, which is converted into acetyl-coenzyme A and further to acetate and ethanol. Our analysis showed that *M. exilis* can obtain glucose from the degradation of glycogen, maltose or starch but can also digest chitin and (1,3)-α-D-glucans present in fungal cell walls and peptidoglycan present in bacterial cell walls, which corroborates its bacterivorous feeding mode (Supplementary Table 2). The biosynthesis of amino acids and nucleotides is limited and dependent on conversion and salvage pathways from compounds acquired from the diet or via transporters. However, *M. exilis* can synthesize dNTPs from NTPs using ribonucleoside-triphosphate reductase (Karnkowska et al., 2019; Novák et al., 2023).

### Bacterial community metabolic capacities

To better understand the metabolic potential of the 24 assembled bacterial genomes, we reconstructed the metabolism of each MAG using EggNOG-mapper and Pathway-Tools. Our results showed a higher number of pathways and reactions compared to *M. exilis*, with considerable diversity in the number of enzymatic reactions, transporters and pathways (from 443 in *Citrobacter portucalensis* to 218 in *Peptostreptococcus russellii*) (Supplementary Table 1). The metabolic maps of the seven most abundant MAGs are provided in Supplementary Figures 2 – 8. The number of reactions and pathways identified in closely related MAGs was very similar, even if the assembly quality differed between MAGs. This suggests that our metabolic analyses were not heavily affected by the level of fragmentation of the MAG assemblies.

We analyzed the roles of community members within the carbon, nitrogen, sulfur, and iron cycles of the community. The results showed that the community likely depends on the oxidation of organic carbon compounds (mainly amino acids and complex carbon compounds), fermentation, and H_2_ generation (Figure 3). The community does not perform methanogenesis, and the produced H_2_ is probably oxidized to water. Our analyses identified *Phascolarctobacterium* sp. as the only community member able to fix N_2_ to ammonium. However, the abundance of this species is low (Figure 2 and Supplementary Table 1) so the N_2_ fixation most likely does not significantly contribute to the ammonium pool. Another potential source of ammonium could be the Dissimilatory Nitrate Reduction to Ammonium (DNRA) pathway (Kaviraj et al., 2024). Components of this pathway, periplasmic nitrate reductase (NapAB) or the nitrate reductase A (NarGHI) and cytochrome c552 nitrite reductase (NrfAH and NrfABCDEF), are present in multiple members of the community (Figure 3, Supplementary Table 3, Supplementary Figures 2, 4 – 7). However, we failed to identify the source of the nitrates and nitrites. These anions are not added to the cultivation medium, and we did not identify enzymes producing nitrate or nitrite either by oxidation of ammonia derived from the amino acid and nucleotide decay or by the biosynthesis of nitrite from aspartate (Sugai et al., 2016). In conformity with the above, the expression of the DNRA pathway in most bacteria, for which the transcriptomic data are available, is extremely low and in the two species, which express it (*B. thetaiotaomicron* and *Parabacteroides* sp.), it is down-regulated in later days (Supplementary Table 3, Supplementary Figures 2, 4 – 7). We conclude that the community does not run a complete nitrogen cycle but depends on the ammonium derived from amino acids and nucleotides supplemented to the medium or originating from dead cells with a small addition through the fixation of nitrogen gas.

**Figure 3:**
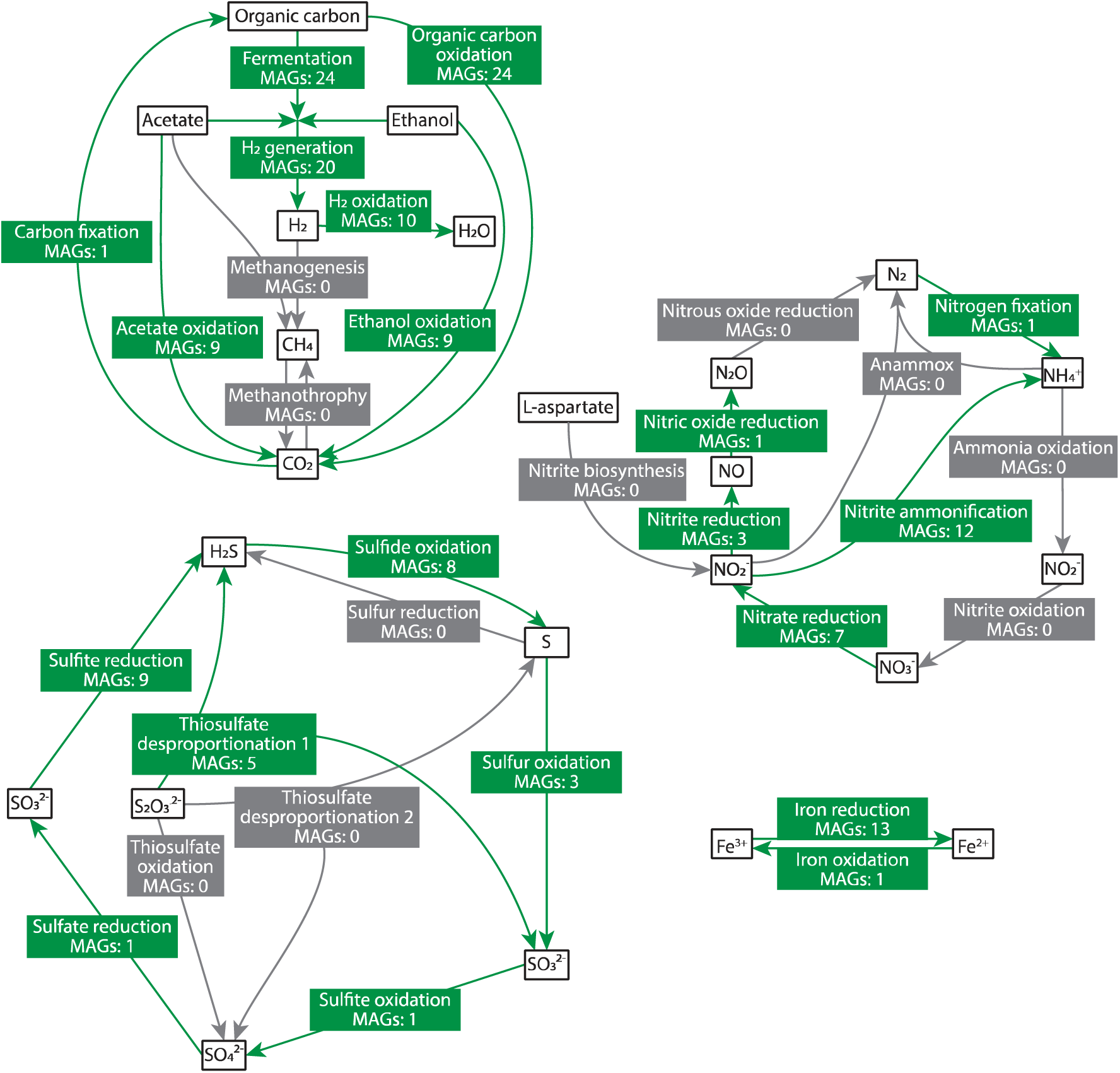
Potential contribution of the bacterial community to the biochemical cycling processes of carbon, nitrogen, iron and sulfur. Labels represent the main steps. Present steps are identified in green. Absent steps are identified in light grey. MAGs: number of MAGs responsible for a step.

In contrast to the carbon and nitrogen cycles, the sulfur cycle is complete. It is maintained by *Desulphovibrio* sp, the only member able to reduce sulfate and oxidize sulfite. We identified eight species that encode the sulfide:quinone oxidoreductase, suggesting that they accumulate sulfur in globules within the cytoplasm (Arieli et al., 1994). The reduction of iron is another important activity in the community, with 13 species involved in the process (Figure 3). In contrast, iron oxidation activity was identified only for *Saezia sanguinis*. (Figure 3).

### The consumption and production of compounds by the community

Triplicates of the cell-free medium were sampled at each time point and their composition was analyzed by non-targeted metabolomics. LC-MS/MS and GCxGC-MS covered a broad range of structural classes of metabolites, detecting 270 distinctive compounds altogether (Supplementary Table 4). Of these, 171 take part in the biological reactions listed on the MetaCyc database (Caspi et al., 2014); hence, they were the only ones used for the follow-up analyses (Supplementary Table 4).

On the day of inoculation (day 0), the medium was already affected by an overnight growth of *C. portucalensis*. Yet it was relatively abundant in nucleotides and their derivates, glyceraldehyde, lactate, fatty acids, alcohols and amino acids, except asparagine, aspartate, cysteine, glutamine and serine, which were below the detection limit at this time point (Supplementary Table 4). On day 2, a drastic evanesce of some compounds was registered, specifically nucleotides and their derivates, most amino acids, glyceraldehyde, glyceric acid, malate, citric acid, and citrulline, suggesting that the community had consumed them. On the contrary, a spectrum of short-chain fatty acids and aromatic compounds putatively derived from the degradation of amino acids, such as tyrosine, phenylalanine and tryptophan, was observed in the media on this day (Supplementary Table 4). The measurements displayed little changes between days 2, 3, and 5, suggesting that the composition of cell-free media stabilized.

We integrated the synthesis and consumption of compounds into an *in silico* metabolic capacity prediction, both for *M. exilis* and the bacterial community, using the Metage2Metabo (m2m) pipeline (Belcour et al., 2020). In these predictions, the culture days were compared pairwise, i.e., day 0 vs. day 2, day 2 vs. day 3 and day 3 vs. day 5. We then classified compounds either as seeds if detected on the first day in pairwise comparison or as targets if they were detected only or their concentration increased on the second sampling day of that pair. This was performed using measured differences in media composition, presence/absence and concentration changes of compounds between sampling days. These results are summarized in Figure 4. From the fact that *M. exilis* represents less than 1.5% of the community metagenomic reads, we infer that its metabolic power is low and the prokaryotic component converts most of the metabolites. Nevertheless, in subsequent analyses, we sought to stress out the metabolic capabilities of this protozoan, which was our main point of interest. Between day 0 and day 2, 137 compounds were classified as seeds and 92 as targets, of which *M. exilis* can synthesize 63 by itself. However, the metabolic capacities of *M. exilis* are not limited to the target compounds, as it can synthesize other important compounds like pyruvate, amino acids, medium-chain fatty acids, and vitamins. An additional 29 target compounds can be synthesized via metabolic interactions of *M. exilis* and the bacterial community (Figure 4). Among these compounds, there are several amino acids. In this pairwise comparison, m2m did not identify any bacteria as an essential member of the community. However, two Bacteroidota bacteria were classified as key members (species occurring in at least one of the minimal community models) (Figure 4). Between day 2 and day 3, we identified 141 compounds as seeds and 72 as targets. According to the m2m pipeline, *M. exilis* can synthesize 68 targets independently and 51 compounds through metabolic interaction with the community (Figure 4). In this comparison, no bacterial species were considered essential or key members of the community. The last comparison, day 3 vs. day 5, showed a similar result. *M. exilis* could synthesize 54 of the 55 compounds identified as targets using 140 compounds as seeds. The interaction between *M. exilis* and the bacterial community allows the synthesis of 49 more compounds and none of the bacteria were considered essential or key members (Figure 4).

**Figure 4:**
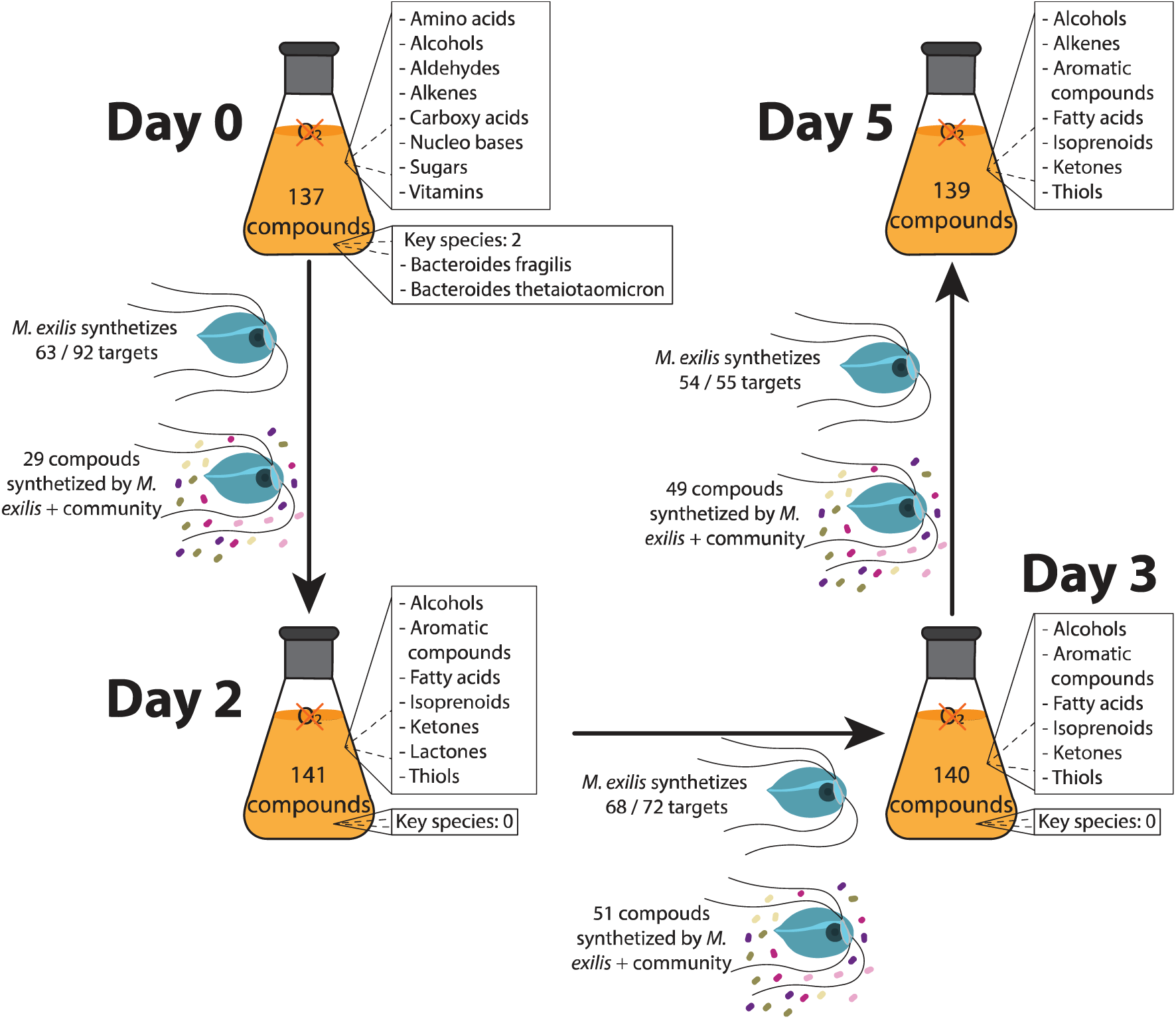
Schematic summary of changes in the culture medium. Chemical classification of significant compounds of identified compounds per day (Supplementary Table 4), number of compounds exhibiting an increase in concentration to the previous day (targets), number of targets that *M. exilis* can synthesize independently, and number of compounds that can be synthesized because of the interaction between *M. exilis* and the bacterial community are shown. Key bacterial species identified by the m2m pipeline are shown for each pairwise comparison.

### *Monocercomonoides exilis* differential expression analysis

To investigate how *M. exilis* metabolism changes during the growth curve, we also compared the expression of transcripts between days 2, 3 and 5, again pairwise. Most significant changes in gene expression were observed between days 2 and 5, while comparisons between consecutive days revealed less significant changes (i.e., day 3 vs. day 2 and day 5 vs. day 3). 2,236 genes were up-regulated and 2,431 down-regulated between days 2 and 5 (Supplementary Table 1), of which the majority, 1703 and 1,3 81 genes, respectively, are annotated as hypothetical; hence their function could not be deciphered.

The changes in expression of all functionally annotated metabolic enzymes and pathways are shown in Supplementary Figure 1 and those discussed below are shown in Figures 5 and 6 in more detail. We observed an increase in the expression of importers of sugars such as α-D-xylose and *sn*-glycerol-3-phosphate towards day 5. Connected to this, there was an increment in the expression of enzymes responsible for the degradation of starch, (1->4)-α-glucan, chitin and other polysaccharides (Figure 5 and Supplementary Table 2). Furthermore, the interconversion between glucose-1-P and glucose-6-P was up-regulated, which may relate to the increased trehalose biosynthesis from glucose-6-P. Trehalose is often used as carbohydrate storage and as a stress protector (Iturriaga et al., 2009). All three pathways for trehalose synthesis were up-regulated on day 5, including the synthesis directly from α-maltose, performed by both α- and β-amylases. The synthesis of glycogen, another storage polysaccharide, was also up-regulated. The expression of most genes involved in carbohydrate metabolism and glycolysis remained unchanged. However, some, such as phosphoenolpyruvate carboxykinase (PEPCK), exhibit down-regulation, suggesting that glycolysis is regulated to produce pyruvate via the up-regulated pyruvate kinase (PK). Later, this pyruvate is converted to acetyl-CoA by pyruvate-ferredoxin oxidoreductase (PFOR) and the up-regulated hydrogenases (HYD) re-oxidize ferredoxins, allowing the cycle to continue. The expression of enzymes of extended glycolysis and ethanol fermentation, i.e., acetate synthase (ACS) producing acetate and aldehyde and alcohol dehydrogenases (ALDH, Adh) producing ethanol, decreases greatly on day 5 for all but one paralogue of ALDH (Supplementary Table 5). ACS1 was consistently the most expressed of these enzymes in all three time points and replicates. Overall, these expression data suggest that acetate and hydrogen are the preferred metabolic end products at all three time points. In contrast to glycolysis, the pentose phosphate pathway may be down-regulated due to the repression of the ribulose-phosphate 3-epimerase (RPE) (Figure 5).

**Figure 5:**
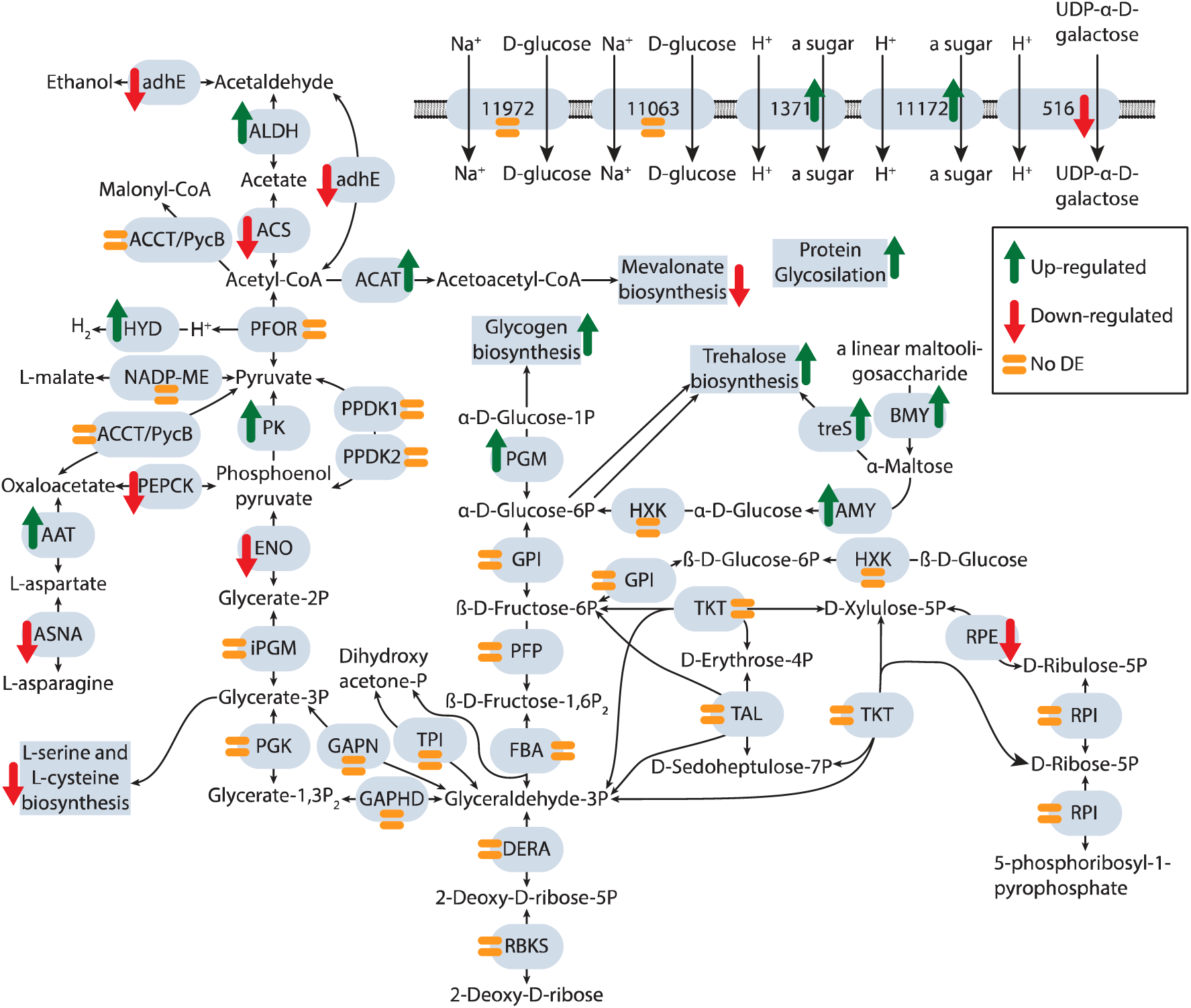
Differential expression of genes involved in carbohydrate metabolism and transport on *Monocercomonoides exilis* between day 5 and day 2. Identified sugar transporters and glycolysis, pentose phosphate pathways, and related reactions (adapted from Karnkowska et al. 2019). Up-regulated genes and processes are marked with a green arrow pointing up. Down-regulated genes and processes are marked with a red arrow pointing down. Non-differentially expressed genes and processes are marked with an orange equal symbol. adhE: Bifunctional acetaldehyde-CoA/alcohol dehydrogenase; ALDH: Aldehyde dehydrogenase; ACS: Acetyl-CoA synthetase (ADP-forming); ACCT/PycB: putative malonyl-CoA:pyruvate transcarboxylase; ACAT: Acetyl-CoA acetyltransferase; PFOR: Pyruvate-ferredoxin oxidoreductase; HYD: [FeFe]-hydrogenase; NADP-ME: NADP-dependent malic enzyme; PK: Pyruvate kinase; PPDK: Pyruvate phosphate dikinase; PEPCK: Phosphoenolpyruvate carboxykinase; AAT: Aspartate aminotransferase; ASNA: Asparagine synthetase; ENO: Enolase; iPGM: 2,3-bisphosphoglycerate independent phosphoglycerate mutase; PGK: Phosphoglycerate kinase; GAPDH: Glyceraldehyde-3-phosphate dehydrogenase; GAPN: NAD(P)-dependent glyceraldehyde-3-phosphate dehydrogenase; TPI: Triose phosphate isomerase; DERA: Deoxyribose-phosphate aldolase; RBKS: Ribokinase; FBA: Fructose-bisphosphate aldolase; PFP: Phosphofructokinase (pyrophosphate-based); GPI: Glucose-6-phosphate isomerase; PGM: Phosphoglucomutase; BMY: β-amylase; AMY: α-amylase; treS: Maltose alpha-D-glucosyltransferase/ alpha-amylase; HXK: Hexokinase; RPDK; ribose-phosphate diphosphokinase; RPI: Ribose-5-phosphate isomerase; RPE: Ribulose-phosphate 3-epimerase; TKT: Transketolase; TAL: Transaldolase.

**Figure 6:**
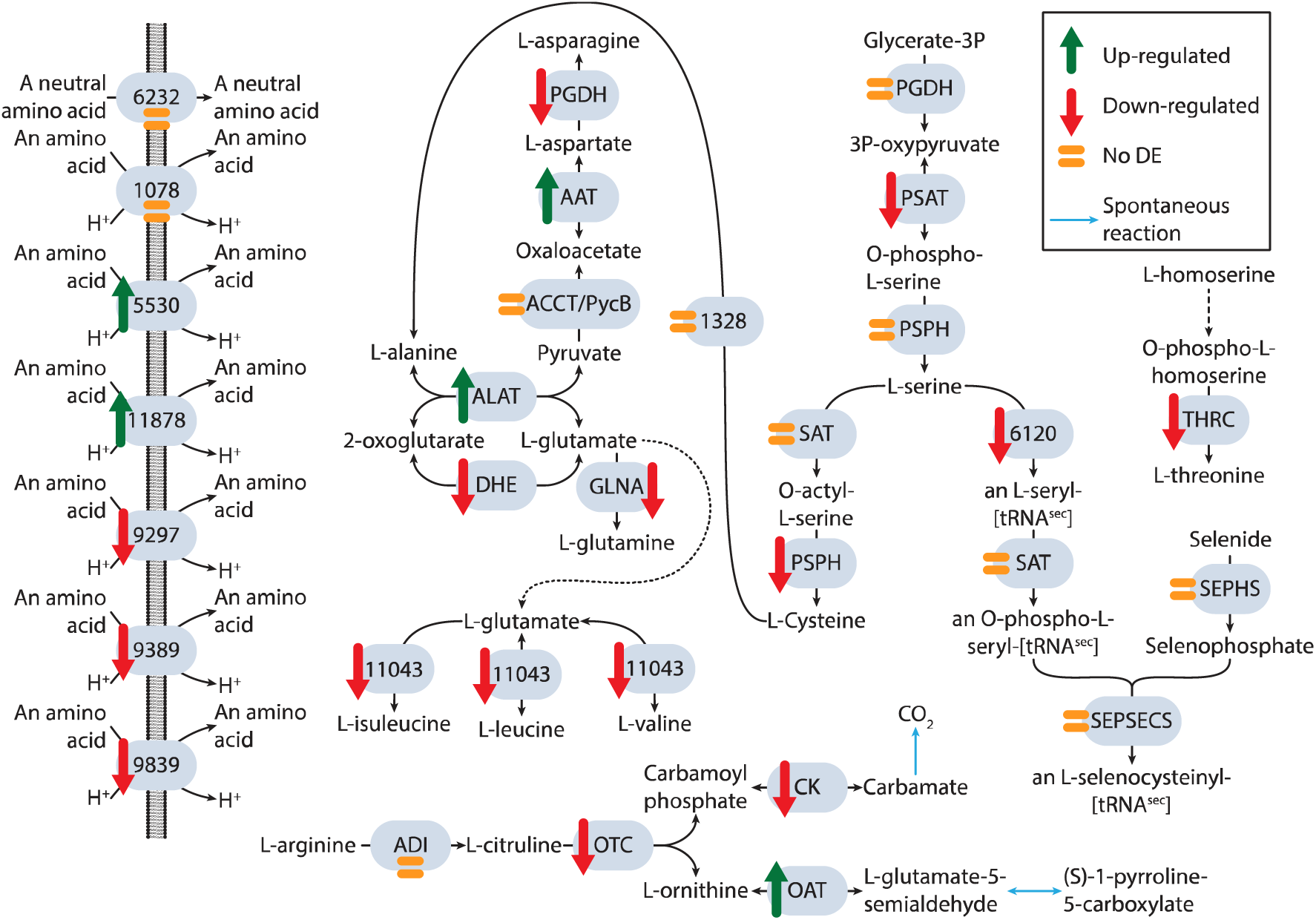
Differential expression of genes involved in amino acid metabolism and transport on *Monocercomonoides exilis* between D5 and D2. Identified amino acid importers, amino acid biosynthesis and arginine catabolism. Up-regulated genes and processes are marked with a green arrow pointing up. Down-regulated genes and processes are marked with a red arrow pointing down. Non-differentially expressed genes and processes are marked with an orange equal symbol. Spontaneous reactions are shown in blue. PGDH: Phosphoglycerate dehydrogenase; PSAT: Phosphoserine transaminase; PSPH: putative phosphoserine phosphatase; SAT: Serine O-acetyltransferase; SEPHS: Selenophosphate synthetase; SEPSECS: O-phospho-L-seryl-tRNA[Sec]:L-selenocysteinyl-tRNA synthase; THRC: Threonine synthase: ASNA: Asparagine synthetase; ALAT: Alanine aminotransferase; ACCT/PycB: putative malonyl-CoA:pyruvate transcarboxylase; AAT: Aspartate aminotransferase; ASNA: Asparagine synthetase; DHE: Glutamate dehydrogenase; GLNA: Glutamine synthetase: ADI: Arginine deiminase: OTC: Ornithine transcarbamylase: CK: Carbamate kinase; OAT: Lysine/Ornithine aminotransferase.

Genes responsible for the phosphorylation of (d)NDP, DNA and RNA polymerization, and protein ubiquitination were up-regulated on day 5 (Supplementary Figure 1). In contrast, nucleoside import, and the salvage pathway of adenosine were down-regulated. Similarly, the mevalonate pathway for the synthesis of isoprenoids from acetoacetyl-CoA was down-regulated (Figure 5 and Supplementary Figure 1). Most amino acid importers displayed lower expression or showed no significant change in expression (Figure 6). This relates to the fact that amino acid content in the media significantly decreased from day 2. Asparagine, serine, and cysteine biosynthesis were down-regulated, as was the ATP-producing arginine deaminase pathway (Figure 6).

Another system that showed significant changes during cultivation was the endomembrane system. Among the proteins involved in the regulation of endomembrane transport, some show significant changes in expression (Supplementary Table 6). The proteins involved in the early secretory pathway show little change, but several proteins involved in the late steps (Law et al., 2022; Shimizu and Uemura, 2022; Tanaka et al., 2017) were significantly up-regulated on day 5 (Supplementary Table 6). These include proteins involved in trans-Golgi-network (TGN) to plasma membrane trafficking (DSCR3A and exocyst complex components Exo84, Sec5 and Sec15, DSCR), multivesicular body formation and sorting (Vsp2, 4a-c, 23, 24a and 60b, charged multivesicular body protein 7), plasma membrane uptake (AP-2a, TSET complex protein TPLATE subunit B), endocytic/digestive pathway (Vps34bc, Vps41b) and endosomes to TGN transport (Vps51b, TRAPP component Bet3 paralogue A, B, syntaxin 6D and 16). On the contrary, four Rab7 paralogues were consistently down-regulated, implying a decrease in lysosome biogenesis on day 5 (Bucci et al., 2000).

At the end of the growth curve, we observed a decline in the number of *M. exilis* cells. Our data showed that most of the enzymes involved in the apoptosis pathway (Teulière et al., 2020) do not show significant expression change (Supplementary Table 7). This suggests that the decline in cell density in later days is not due to programmed cell death but most likely due to uncontrolled decay combined with slow growth caused by nutrient deficiencies.

### Bacterial community differential expression analysis

Our data only allowed to perform differential expression analysis for the seven most abundant bacteria: *B. fragilis*, *B. thetaiotaomicron*, *B. intestinalis*, *Parabacteroides* sp., *K. gyiorum*, *Fusobacterium varium* and *C. portucalensis*. These species represent 80-90% of the community DNA reads, a proxy measure of their abundance. These bacteria showed a variable number of up- or down-regulated genes (Supplementary Table 1); for example, only 9% of *B*. intestinalis. transcripts changed in expression between day 2 vs. day 5, while in *Parabacteroides* sp., it was 51%. Changes in the expression of metabolic enzymes and pathways of these bacteria are indicated in Supplementary Figures 2 – 8.

We focused specifically on pathways absent in *C. portucalensis* but present in other members of the community, as these may explain why we failed to grow *M. exilis* with only *C. portucalensis* as food. First*, F. varium* and *Tissierella* sp. can ferment lysine to acetate, butanoate and ammonia, while in *C. portucalensis*, this pathway is incomplete. Furthermore, *B. intestinalis, S. sanguinis, K. gyiorum, Phascolarctobacterium* sp. and *Tissierella* sp. encode genes for the synthesis of the essential polyamine spermidine and the phospholipid phosphatidylcholine. While phosphatidylcholine is one of the major components of cell membranes (Zhukov and Popov, 2023), spermidine is involved in an array of biological processes, such as maintenance of membrane potential, control of intracellular pH, lipid metabolism and regulation of cell growth, proliferation, and death (Michael, 2016). In both cases, our metabolic analysis fails to identify a pathway that can synthesize these compounds *de novo* in *M. exilis*, suggesting that the eukaryote may depend on the uptake of these compounds from these prokaryotes.

Iron uptake is another relevant function for which no pathway is known in *M. exilis*. Several bacteria encode for transporters of Fe^2+^, the anaerobic Feo import system, the aerobic Efe system and the Sit system (Braun and Hantke, 2013) (Supplementary Table 8). Our data showed that in all bacteria with expression data, only the Sit system in *K. gyiorum* is up-regulated between day 2 and day 5. In the rest of the bacteria, all the Fe^+2^ transporters are down-regulated or not significantly differentially expressed between these days (Supplementary Figures 2 – 8). Of importance is also another type of molecules used for iron chelation in bacteria known as siderophores. These iron-chelating molecules are secreted into the environment and their iron-bound forms are subsequently selectively imported (Kramer et al., 2020). Our data showed that some, but not all, community members produce siderophores, *C. portucalensis* and *S*. *sanguinis* synthesize enterobactin, while *F. varium* synthesizes staphylopine, and *K. gyiorum* a periplasmic iron-binding protein. Interestingly, all bacteria, but not *M. exilis*, encode transport systems for siderophores or transferrin. Of these, we identified components of the Ton uptake system (TonB, ExbB and ExbD) (Braun and Hantke, 2013) in most gram negative bacteria except for ExbB absent from *Morganella morganii* (Supplementary Table 8). Consistently, we identified several TonB-dependent receptors and transporters, namely the enterobactin transporters FepA and FbpABC and the transferrin receptor TbpA in many members of the community. The presence of these proteins and protein complexes makes the transport of siderophores or heme groups across the outer membrane possible (Braun and Hantke, 2013). Our data showed that the Ton uptake system and the receptor TbpA are up-regulated in *F. varium, K. gyiorum* and members of the Bacteroidota in later days of the culture (Supplementary Figures 3 – 5, 7,8). The described pattern of presence and absence suggests that the bacterial community contains only a few species able to synthesize siderophores and all the other members are “cheaters” that uptake siderophores produced by others (Butaite et al., 2017; Leventhal et al., 2019). From this point of view, the siderophores’ producers are key players in the prokaryotic community but do not facilitate the uptake of iron into *M. exilis.* Once iron is imported into the cell, it cannot be kept free in the cytoplasm. One solution would be its coordination within different kinds of porphyrins (Bryant et al., 2020; Dailey et al., 2017; Layer, 2021). We observed two main porphyrins, siroheme and heme *b*. Heme *b* can be synthesized aerobically or in an oxygen-independent manner, while siroheme is synthesized only independently of oxygen (Dailey et al., 2017; Layer, 2021). Our data showed that *C. portucalensis* can synthesize both compounds. Still, under anaerobic conditions, the synthesis of heme *b* is down-regulated, while that of siroheme is up-regulated on day 5 (Supplementary Figure 2). The opposite situation was observed in *Bacteroides fragilis,* where oxygen-independent synthesis of heme *b* was up-regulated, yet siroheme synthesis underwent down-regulation (Supplementary Figure 4). Some bacteria in the community can synthesize heme *b* from siroheme. However, we lacked transcriptome data for their assessment due to their low abundance.

## Discussion

The only study of chinchilla gut microbiota composition (O’ Donnell et al., 2017) reports Betaproteobacteria as the dominant phylum and *Parabacteroides* and *Barnesiella* as the dominant genera. Our study of an *M. exilis* culture derived from the gut of chinchilla identified 30 bacterial species, of which *Fusobacterium varium* was the most abundant at the beginning of the experiment, steadily replaced by the Bacteroidetes group, represented by *Bacteroides fragilis*, *Parabacteroides* sp. and *B. thetaiotaomicron* on day 5. *Parabacteroides* sp. was the second most abundant bacteria on day 5, yet we failed to detect any *Barnesiella* species. Only two Betaproteobacteria species were identified in our study, but they are not among the most abundant species. At the level of bacterial phyla, Bacteroidota and Bacillota are the most abundant in our culture, which, on a broad scale, resemble the composition of the gut microbiota of Guinea pigs (Crowley et al., 2017). These comparisons indicate similar compositions of the prokaryotic community between the gut of caviomorph rodents and our *M. exilis* culture, yet the culture community has changed due to long-term cultivation.

Given that *M. exilis* represents around 1% of the read percentage, the biochemical processes in the community are driven by prokaryotes. Of the three main cycles, only the sulfur cycle seems to be complete and contains reactions converting all basic inorganic sulfur compounds (Figure 3). Our analyses infer that the community dominantly oxidizes organic carbon substrates, performs fermentation and generates hydrogen, which is further oxidized to water (Figure 3). Since the ability to fix carbon from CO_2_ is present only in *Eubacterium maltosivorans,* the least abundant species, the community depends on the supply of organic carbon. This is consistent with our observation that organic carbon sources (glyceraldehyde, lactate, fatty acids, alcohols and most amino acids) present on day 0 are quickly consumed.

The community does not run a complete nitrogen cycle. While there is a small level of N_2_ fixation due to the contribution of *Phascolarctobacterium* sp., this bacterium is among the less abundant members of the community, and most likely, its N_2_ fixation does not cover the demands for ammonium. Another pathway for ammonium synthesis, the DNRA pathway using the periplasmic nitrate reductase or the nitrate reductase A, and the membrane-bound nitrate reductase nrf, is encoded by some bacterial species (Kaviraj et al., 2024). This pathway uses nitrate as electron acceptor and allows building membrane potential (Cabello et al., 2019). However, in all the bacteria with transcriptomic data, the DNRA is not expressed or downregulated in later days. The nitrogen sources for building a biomass thus are amino acids and nucleobases, which are components of the growth medium and are measurable on the day 0 with a small supplement of ammonia synthesized by *Phascolarctobacterium* sp.

The metabolomic analyses of the soluble media clearly showed that on day 2, when the population of *M. exilis* only began to grow, essentially all soluble carbon sources, nucleotides and most amino acids had been already consumed (Supplementary Table 4). The fact that *M. exilis* maintains active growth after day 2 strongly suggests that the protist uses bacterial prey as the major source of carbon, nitrogen, and energy to build its biomass. A putative exception could be arginine, which was detected in later days and may serve as a potential substrate for energy metabolism (Schofield et al., 1992; Yarlett et al., 1996). Grazing on bacteria has been previously documented by electron microscopy (Treitli et al., 2018) and the fact that the protist encodes lysozymes amongst a broad array of glycosyl hydrolases (Supplementary Table 2) also provides molecular tools for feeding on bacteria. There is no consistent pattern that would indicate up- or down-regulation of lysozymes as the culture ages. The protist, however, displays a significant up-regulation of several genes involved in the late pathways of the endomembrane system on day 5. At the same time expression of many genes encoding for proteins regulating these processes remained unchanged and, notably, four paralogues of Rab7 small GTPase, known for its role in lysosome activation (Bucci et al., 2000), were significantly down-regulated on day 5 (Supplementary Table 6). We interpret this pattern as an indication of up-regulation of exocytosis and secretion rather than bacterial feeding, which is also consistent with the decrease in *M. exilis* cell counts at this stage of the culture.

Most *M. exilis* genes that exhibit differential expression completely lack functional annotation, providing us with a rather large transcriptomic dark matter consisting of approx. three thousand genes that may convey some functional and adaptive value to the protist, but we know absolutely nothing about it. Among the known metabolic genes, we observed an up-regulation of the importers of sugars, α-D-xylose and *sn*-glycerol-3-phosphate, which is likely the reaction of the cell to the absence of these substrates. Another functional category of genes up-regulated on day 5 is the enzymes responsible for starch degradation and synthesis of trehalose and glycogen, indicating the onset of starvation. The degradation of nucleosides and nucleotides and the biosynthesis of 5-phospho-α-D-ribose 1-diphosphate (PRPP) are up-regulated, suggesting an increment in the storage of nucleotides and phosphate donors. On the other hand, glycolysis, extended glycolysis, fermentation, pentose phosphate pathway, amino acids biosynthesis, arginine deaminase pathway and mevalonate pathway involved in isoprenoid biosynthesis are generally down-regulated on day 5, with few exceptions. The same holds for the synthesis of very long and ultra-long fatty acids and the aminoacyl-tRNA synthetases, indicating the decline of proteosynthesis.

Although we hoped to find some metabolic dependencies between *M. exilis* and prokaryotes, potentially explaining the inability to grow the flagellate axenically using nutrients dissolved in the medium, we did not reveal any clear link. This obviously does not mean that some have remained unobserved or unnoticed, as the amount and complexity of the data are significant. The *in silico* metabolic capacity prediction by Metage2Metabo (Figure 4) has shown that the compounds detected in the media at the time points can be synthesized by the reactions and pathways found in the community. Importantly, none of the community members has been found to be essential at any time point, though some may provide key compounds for others, e.g., siderophores. This suggests that the community is based on broad redundant foundations, which certainly contributes to its long-term stability. However, we infer that carbon and nitrogen could be limited factors that can disturb this tendency.

## Material and methods

### Cell culturing

700 mL of TYSGM-9 medium in 2x 1 L glass bottles were inoculated with 500 µL of *Citrobacter portucalensis* culture before the inoculation of the whole community. This strain of *C. portucalensis* was previously isolated from the original culture during the isolation of *M. exilis*. After inoculation, the flasks were closed and allowed to grow standing at 37°C. The next day, the two flasks were merged in a 2 L bottle and mixed thoroughly. 45 mL of culture were removed and used to isolate DNA, RNA and metabolomic analyses of day 0. The rest of the medium was inoculated with approx. 125 mL of *M. exilis* culture (2.5×10^5^ cells.mL^-1^). After inoculation, the media was thoroughly mixed and divided into 50-mL tubes containing 45 mL of media, closed, and incubated at 37°C. Each day, three tubes were removed, one for each replicate. From each tube, 20 mL of culture were used for DNA isolation, 20 mL for RNA isolation and 5 mL for metabolomics. Before each nucleic acid isolation, *M. exilis* cells were counted using a Neubauer cytometer.

### DNA isolation

All DNA isolations were performed using the Dneasy Blood and Tissue Kit (Qiagen). Each culture was centrifuged at 1500 x*rcf* for 10 min at 4°C. The supernatant was separated into a fresh 50 mL tube and centrifuged again at 6000 x*rcf* for 10 min at 4°C. The pellet from the first centrifugation was resuspended in 200 µL of PBS and DNA was isolated using the cultured cells protocol of the Dneasy Blood and Tissue Kit (Qiagen). The second pellet, obtained after centrifugation at 6000 x*rcf*, was resuspended in 200 µL of PBS and DNA was isolated using the gram-positive protocol of the Dneasy Blood and Tissue Kit (Qiagen). For each isolation, the DNA was eluted in 100 µL of elution buffer. The DNA from the two pellets was merged into one DNA sample. The quality of the DNA was estimated using Nanodrop and the concentration was measured using the QuantiFluor ONE dsDNA System (Promega).

### RNA isolation

Each culture was centrifuged at 1500 x*rcf* for 10 min at 4°C. The supernatant was separated into a fresh 50 mL tube and centrifuged again at 6000 x*rcf* for 10 min at 4°C. Both pellets were resuspended together in 1 mL Tri-Reagent (Sigma-Aldrich) and the total RNA was isolated according to the manufacturer’s procedure. The obtained RNA pellet was resuspended in 60 µL of RNA-grade H_2_O and heated at 37°C for 10 min to allow full resuspension.

The total RNA was further DNAse-treated. For this purpose, we added 6 µL of rDNAse buffer and 0.6 µL rDNAse (Macherey-Nagel) per sample and incubated at 37°C for 10 min. Samples were subsequently re-isolated with Tri-Reagent (Sigma-Aldrich) as before and finally resuspended in 30 µL of RNA-grade H_2_O. RNA concentration was measured using Nanodrop.

### Identification of compounds within the growth medium

For metabolomics, cells were pelleted as described for DNA and RNA isolation (see above). The supernatant obtained in the second centrifugation was filtered (Whatman polycarbonate filter, 3 µm pore size)-sterilized and stored at −80°C. Metabolites released to the media were determined using a non-targeted metabolomics approach. To capture both volatiles and non-volatiles, a combination of two techniques was employed – Liquid Chromatography analysis in connection with Orbitrap Fusion mass spectrometer (LC-MS/MS; Orbitrap Fusion, Q-OT-qIT, Thermo Scientific) and two-dimensional comprehensive gas chromatography in connection with mass spectrometer (GCxGCTOF-MS; Pegasus 4D, Leco Corp.).

For LC-MS/MS, media samples harvested in triplicates were precipitated with acetonitrile in a ratio of 4:1 (ACN:media) and centrifugated for 20 min at 16000 x*rcf* at 5=C. 100 µL of each supernatant was collected, fully evaporated and dissolved in 50 µL 10% (v/v) ACN. 15 µL were injected into a ProntoSIL column, 150 x 3.0 mm, 3 µm (Bischoff Chromatography). Compounds were eluted under the following conditions: 0 - 3 min 100% A, 3 - 18 min linear gradient to 100% B, 18 - 20 min 100% B using mobile phases. A: 2% (v/v) ACN, 10 mM HCOONH_4_, 0.1% (v/v) FA; B: 99% (v/v) ACN, 0.1% (v/v) FA with a flow of 0.4 mL.min^-1^. The separated compounds were ionized in an electrospray ion source in positive polarity mode. Master scans of precursors from 80 to 1000 m/z were performed in Orbitrap at 120 K resolution with an intensity threshold filter 2.0 × 10^4^. Tandem MS was performed by isolation in the quadrupole, HCD fragmentation with a stepped collision energy of 15, 30, and 45% and an isolation window of 1.6 m/z. The MS2 ions were analysed and detected in Orbitrap with a set resolution of 30k and a max injection time of 300 ms. The dynamic exclusion duration occurred every 20 s with 10 ppm tolerance around the selected precursor. The data processing, including chromatographic peaks alignment, was performed using Compound Discoverer 3.3 (Thermo Scientific). Quality control samples were employed for the data correction. The mzCloud libraries were used as a tool for a fragmentation spectra comparison and the compound structure assignment.

Volatiles were analyzed using GCxGC-MS and a headspace solid phase microextraction (HS-SPME) on fiber (DVB/CAR/PDMS_grey; Supelco, USA). A volume of 1 mL of media samples was placed in a 20 mL glass vial and 10 µL of an internal standard (2,2,2-trifluoro ethanol, Fluka, 0.11 mg.mL^-1^) was added. Samples were incubated for 10 min at 40°C before extraction. The extraction was carried out for 5 minutes. The volatiles were analyzed using a combination of polar and mid-polar separation columns for the separation (primary column: Stabilwax-DA (30 m x 0.25 mm, Restek, USA); secondary column BPX-50 (1.38 m x 0.1 mm, SGE, Australia)). Other parameters were set as follows: inlet temperature 220°C, split 5 modes, constant He flow 1 mL.min^-1^, modulation time 3 s (hot pulse 0.7 s), modulation temperature offset with respect to the secondary oven 20°C. The temperature program applied on the primary oven was 30°C (hold 1.5 min), then increased to 110°C (8°C.min^-1^), followed by an increase to 250°C (25°C.min^-1^) to 250°C (hold 5 min). The temperature offset applied on the secondary column was +5°C. Transferline temperature was held at 280°C. The mass spectrometer was equipped with an Electron Ionization ion source (energy of 70eV was applied),and Time-of-Flight analyser enabling a united mass resolution. The scanned mass range was 29 – 400 *m/z*. The ion source chamber was held at 250°C.

To extend the range of compounds analyzable by gas chromatography to polar non-volatiles, the media samples underwent two consecutive derivatization procedures – oximation (25 mg.mL^-1^ of methoxyamine hydrochloride in anhydrous pyridine hydrochloride, Sigma-Aldrich) and silylation. Two different agents were used for silylation: N,O-Bis(trimethylsilyl)trifluoroacetamide with trimethylchlorosilane (BSTFA + 1% (v/v)TMSC) and N-tert-Butyldimethylsilyl-N-methyl trifluoroacetamide (MTBSTFA), both from Sigma-Aldrich.

For silylation using BSTFA + 1% (v/v) TMSC, 50 µL were transferred to a 2 mL glass vial and 2 μL of internal standard were added (Adonitol, 424 μL.mL^-1^). The liquid was dried in a vacuum concentrator and reconstituted back by adding 30 μL of anhydrous pyridine and 30 μL of an oximation agent. The samples were then incubated in a Thermomixer (Eppendorf) at 40°C for 2 hours at constant shaking (1500 rpm), after which 40 μL of neat solution of BSTFA + 1% (v/v) TMSC mixture were added to the samples and were further incubated at 70°C for 30 min at constant shaking (1500 rpm). Before analysis, 400 μL of hexane was added to the sample. For silylation using MTBSTFA, the derivatization process is essentially the same as the derivatization using BSTFA + TMSC except that 30 μL of media sample are used, plus 5 μL of nor-Valine (550 μL.mL^-1^) as an internal standard and 20 μL of oximation agent.

Metabolites were analyzed as oximated trimethylsilylated or tert-butyl silylated derivatives. A combination of nonpolar and mid-polar separation columns was used for the separation; a primary Rxi-5SIL MS column (30 m x 0.25 mm, Restek, USA) and a secondary BPX-50 column (1.39 m x 0.1 mm, SGE, Australia). Other parameters were set as follows: inlet temperature 290°C, splitless mode, constant He flow 1 mL.min^-1^, modulation time 3s (hot pulse 0.7s), modulation temperature offset with respect to the secondary oven 15°C. The temperature program applied to the primary oven was 50°C (hold 1 min), then increased (8°C.min^-1^) to 320°C (hold 2 min). The temperature offset applied on the secondary column was +5°C. Transferline temperature was held on 280°C. The scanned mass range was 85 – 800 *m/z*. The ion source chamber was held at 280°C. Data were processed in ChromaTOF v4.5 software. Detected analytes were quantified relatively after normalization to the internal standards. Metabolites detected by all the methods used were identified by comparison of their mass spectra with those available in the main NIST mass library, Fiehńs mass library of silylated compounds, and in-house-built libraries. The retention index was determined using linear hydrocarbons. In the cases where the same compounds were detected in both derivatizations, the highest signal was taken to the final table.

### Library preparation and sequencing

For the DNA samples, sequencing libraries were prepared from 1 µg of gDNA using the Illumina TruSeq DNA PCR free (Illumina) library preparation kit according to the manufacturer’s protocol. The prepared libraries were sequenced on an Illumina HiSeq 4000 with 2×150 bp reads.

For RNA, 1 µg of total RNA per sample was used for library preparation. Ribosomal RNA depletion was performed using the NEBNext rRNA Depletion Kit for Bacteria (New England Biolabs), spiked with a custom oligo pool. The oligo pool was designed based on bacterial and eukaryotic ribosomal sequences that were identified from previous sequencing projects of *M. exilis* (Karnkowska et al., 2016; Treitli et al., 2021). After rRNA depletion, libraries were prepared using the NEBNext® Ultra™ Directional RNA Library Prep Kit (New England Biolabs) according to the manufacturer’s protocol. The prepared libraries were sequenced using Illumina NextSeq 500 with 2×75bp reads.

### Metagenome assembly and binning

The metagenomics raw reads were quality-checked with FASTQC v0.11.5 and trimmed using Trimmonatic v0.39 (Bolger et al., 2014) (ILLUMINACLIP:TruSeq3-PE-2.fa:2:30:10, LEADING:20, TRAILING:20, SLIDINGWINDOW:5:20 MINLEN:50). To reduce complexity during assembly, the eukaryotic reads were removed by mapping the reads against the *M. exilis* genome (Treitli et al., 2021) using BBMap (minidentity=0.98, idfilter=0.98) included in BBTools v38.90 (https://sourceforge.net/projects/bbmap/) and taking only unmapped reads.

The filtered reads were assembled independently for each sample using metaSPAdes v3.14.0 (Nurk et al., 2017) with k-mers 21,33,55,77,99,127. A reiterative process was used for binning metagenome-assembled genomes (MAGs) within the samples. First, contigs above 2kbp were binned using MaxBin v2.2.7 (Wu et al., 2016) (minimum probability 0.9, marker set 107) and tetraESOM (Dick et al., 2009). Then, bins were manually checked based on the markers identified by MaxBin using Blastp against nr database (April 2021). For each potential MAG, rRNA genes were identified using Prokka v1.14.6 automatic annotation pipeline (Seemann, 2014) and used as controls together with the marker genes. Lastly, reads corresponding to trustful bins (markers and rRNA pointing to the same species) were removed from the data set by mapping the reads back to the identified bins using BBSplit (minidentity=0.98, idfilter=0.98) included in BBTools v38.90. This process was iterated and stopped when no new clean MAG was created. In total, 24 prokaryotic MAGs were assembled. The contribution of each prokaryotic MAG and *M. exilis* to the whole community per sample was calculated as the percentage of reads mapped to each genome assembly.

To identify possible viral particles and plasmids in the culture, the filtered reads were assembled independently for each sample using metaviralSPAdes (Antipov et al., 2020) and metaplasmidSPAdes (Antipov et al., 2019), respectively, with k-mers 21,33,55,77,99,127. Candidates contigs were verified using viralVerify tool (github.com/ablab/viralVerify.git) and Prokka v1.14.6.

### Identification of prokaryotic MAGs

The taxonomic classification of the 24 prokaryotic MAGs was computed using classify_wf, included in GTDB-Tk (Chaumeil et al., 2022; Eddy, 2011; Harris et al., 2020; Hyatt et al., 2010; Matsen et al., 2010; Ondov et al., 2016; Parks et al., 2020; Price et al., 2010; Shaw and Yu, 2023; Sukumaran and Holder, 2010) with standard parameters. The completeness and contamination of every MAG were estimated based on the presence/absence and number of copies of the 107 marker genes used by MaxBin (see above).

### Annotation and metabolic capacities of prokaryotic MAGs

The annotation of the 24 prokaryotic MAGs was performed using Prokka v1.14.6. The functional annotation of each MAG was made using EggNOG-mapper v1.0.3-35 on emapper DB 2.0 (Cantalapiedra et al., 2021; Huerta-Cepas et al., 2019) (standard parameter except minimum % of query coverage: 50 and minimum % of subject coverage: 50). EggNOG-mapper predictions and Prokka annotations were combined using emapper2gbk, included in the Metage2Metabo pipeline (Belcour et al., 2020), to create a compatible input for Pathway-Tools v24.5 (Karp et al., 2020).

A metabolic database was created for each MAG using Pathologic (Karp et al., 2011), included in Pathway-Tools (standard parameters except name matching that was turned off). All databases were manually curated using the Assign Probable Enzymes, Transport Inference Parser and Rescore Pathways tools included in Pathologic. Predicted pathways were manually checked to remove highly incomplete pathways.

### Identification of metabolic capacities of *Monocercomonoides exilis*

The genome of *M. exilis* was downloaded from NCBI. The metabolic capacities of this species were predicted using Pathologic, included in Pathway-Tools, with standard parameters. The resulting database was manually curated using the Assign Probable Enzymes, Transport Inference Parser and Rescore Pathways tools included in Pathologic. Information from previous curations (Karnkowska et al., 2019, 2016; Treitli et al., 2021) and EggNOG-mapper was included at this step.

Carbohydrate-metabolizing enzymes were identified by computing searches against CAZy databases using METABOLIC-G with default parameters (Zhou et al., 2022).

### Transcriptome processing and identification of differentially expressed genes

Meta-transcriptomic raw reads were quality-checked with FASTQC v0.11.5 and trimmed using Trimmonatic v0.39 (ILLUMINACLIP:TruSeq3-PE-2.fa:2:30:15, LEADING:15, TRAILING:15, SLIDINGWINDOW:4:15 MINLEN:50), resulting in 47 million read pairs per sample on average (except for sample D5R3 where the number of read pairs was 77 million). Reads were mapped and the expression of transcripts of prokaryotes and *M. exilis* was quantified using Salmon (Patro et al., 2017) with default parameters.

Due to the low mapping rate and a low number of reads, the differential expression was assessed only for *M. exilis* and the seven most abundant prokaryotic genomes: *Bacteroides fragilis, Bacteroides thetaiotaomicron, Bacteroides intestinalis*, *Parabacteroides* sp.*, Kerstersia gyiorum, Fusobacterium varium* and *Citrobacter portucalensis*. Raw reads were re-mapped onto the eight selected genomes using Bowtie2 (Langmead and Salzberg, 2012). Reads mapping to protein-coding sequences were counted with the FeatureCounts program of the subread package v2.0.3 (Liao et al., 2014). Read counts were normalized using the DESeq2 package v1.32.0 (Love et al., 2014) in R v4.1.1 (R Core Team, 2021). Normalisation was based on taxon-specific scaling (Christel et al., 2018; Klingenberg and Meinicke, 2017).

### Identification of the major biochemical cycles in the community

The metabolic capacities of the prokaryotic community were also analyzed using METABOLIC-C (Zhou et al., 2022) with default parameters, except tax genus, to understand their roles within the community’s major biochemical cycling. We manually verified the results concerning the carbon, nitrogen, sulfur, and iron cycle by combining the results from METABOLIC-C and EggNOG-mapper from previous analyses.

### Identification of the metabolic relationships between *M. exilis* and the prokaryotic community

All identified reactions in *M. exilis* and the 24 prokaryotic MAGs metabolic database were exported as SBML format from Pathway-Tools. The Metacom tool implemented in the metage2metabo (m2m) pipeline was used to identify metabolic interactions between *M. exilis* and the rest of the community. In this analysis, the sampling days were compared pairwise, with *M. exilis* defined as the host. In each pairwise comparison, compounds that could be identified in the first sampling point were defined as seeds. Meanwhile, those whose concentration increased in the second sampling point were defined as targets.

## Acknowledgment

We thank Professor Joel Dacks for his advice on the analysis of the differentially expressed protein related to the endomembrane system. This project has received funding from the European Research Council (ERC) under the European Union’s Horizon 2020 research and innovation program (grant agreement No. 771592 to VH) and the Centre for Research of Pathogenicity and Virulence of Parasites (registration no. CZ.02.1.01/0.0/0.0/16_019/0000759). The metagenome assemblies were computed using the resources provided by the e-INFRA CZ project (ID:90254), supported by the Ministry of Education, Youth and Sports of the Czech Republic.

## Data Availability Statement

DNA reads, RNA reads and the complete 24 MAGs are available under a BioProject with accession number PRJEB70623. MAGs are available with accession numbers from ERS17699982 to ERS17699999.

