## Supplementary Figure for "Comprehensive analysis of the microbial consortium in the culture of flagellate *Monocercomonoides exilis*"

### Supplementary Figure legends

**Supplementary Figures 1 - 8** Metabolic cellular overview of 1: *Monocercomonoides exilis*; 2: *Citrobacter portucalensis*; 3: *Fusobacterium varium*; 4: *Bacteroides fragilis*; 5 *Bacteroides thetaiotaomicron*; 6 *Bacteroides intestinalis*; 7 *Parabacteroides* sp.; 8 *Kerstersia gyiorum*. For each reaction, the E.C. number and the assigned gene are shown. Transcriptomic data was mapped for the different genes. Up-regulated genes are colored in green. Down-regulated genes are colored in red. No differentially expressed genes are colored in orange.

### Supplementary Table legends

**Supplementary Table 1** Identified MAGs and *Monocercomonoides exilis* assembly statistics; relative read abundance in percentage; Metabolic prediction statistics; Number of differentially expressed genes (number of hypothetical genes for *M. exilis* are indicated in brackets).

**Supplementary Table 2** Carbohydrates-active enzymes identified in *M. exilis* using METABOLIC-G. For each enzyme, the CAZyme family, function and differential expression are indicated.

**Supplementary Table 3** Identified genes involve in the dissimilatory nitrate reduction to ammonium (DNRA) pathway in the bacterial community. Read counts per kb of genes involved in this pathway.

**Supplementary Table 4** Compounds identified in every sample point. Compounds that take part in any biological reaction listed on the MetaCyc database are marked in bold.

**Supplementary Table 5** Read counts per kb of enzymes of extended glycolysis

**Supplementary Table 6** Differential expression of the endomembrane system genes in *M. exilis*. Up-regulated genes are marked with a "+", down-regulated genes are marked with a "-" and non-differentially expressed genes are marked with a "0".

**Supplementary Table 7** Differential expression of the apoptosis pathway genes in *M. exilis*. Up-regulated genes are marked with a "+", down-regulated genes are marked with a "-" and non-differentially expressed genes are marked with a "0".

**Supplementary Table 8** Identified genes involved in the uptake of $Fe^{2+}$, heme groups and siderophores

### *Monocercomonoides exilis* Cellular Overview D5 vs. D2

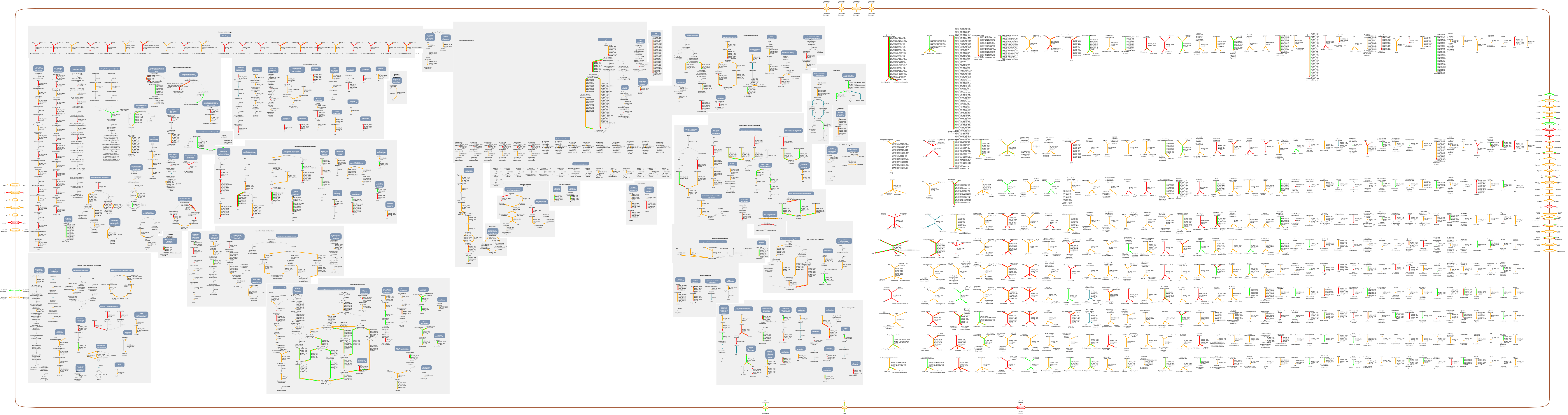

This diagram was automatically generated by SRI International Pathway Tools version 25.5, authors S. Paley and P.D. Karp.

### *Citrobacter freundii* Cellular Overview D5 vs. D2

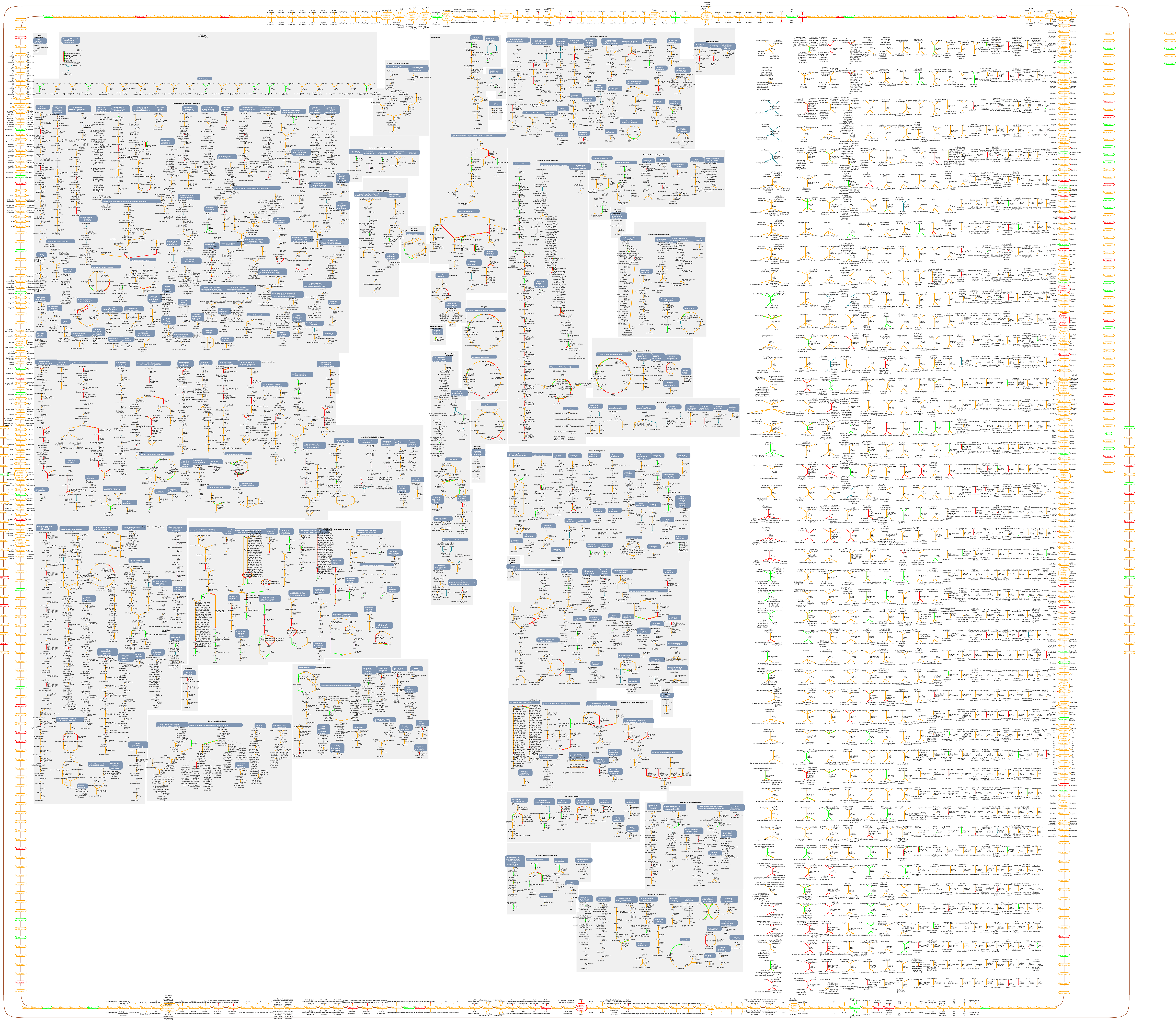

This diagram was automatically generated by SRI International Pathway Tools version 25.5, authors S. Paley and P.D. Karp.

### *Fusobacterium varium* Cellular Overview D5 vs. D2

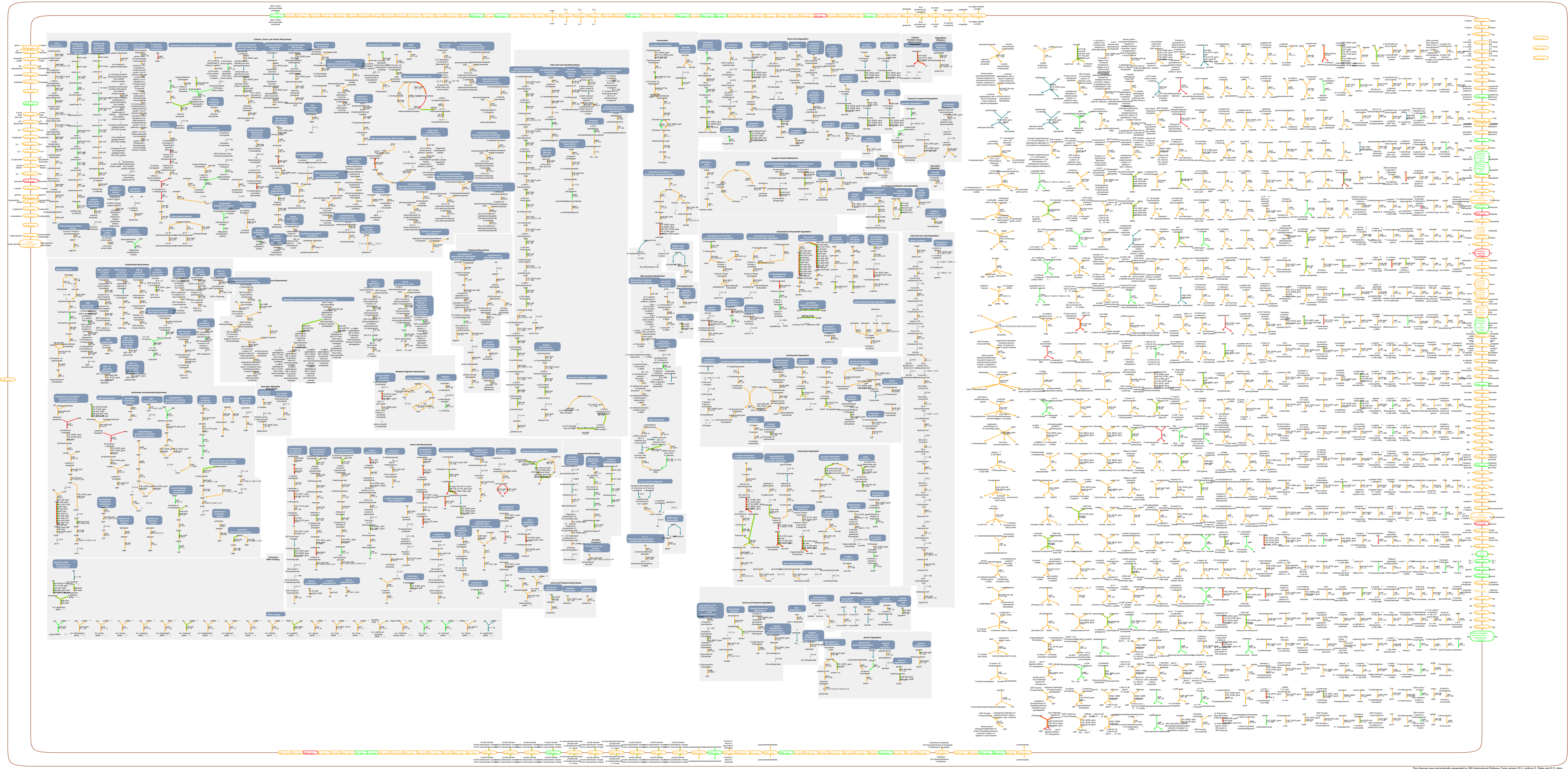

This diagram was automatically generated by SRI International Pathway Tools version 25.5, authors S. Paley and P.D. Karp.

### *Bacteroides fragilis* Cellular Overview D5 vs. D2

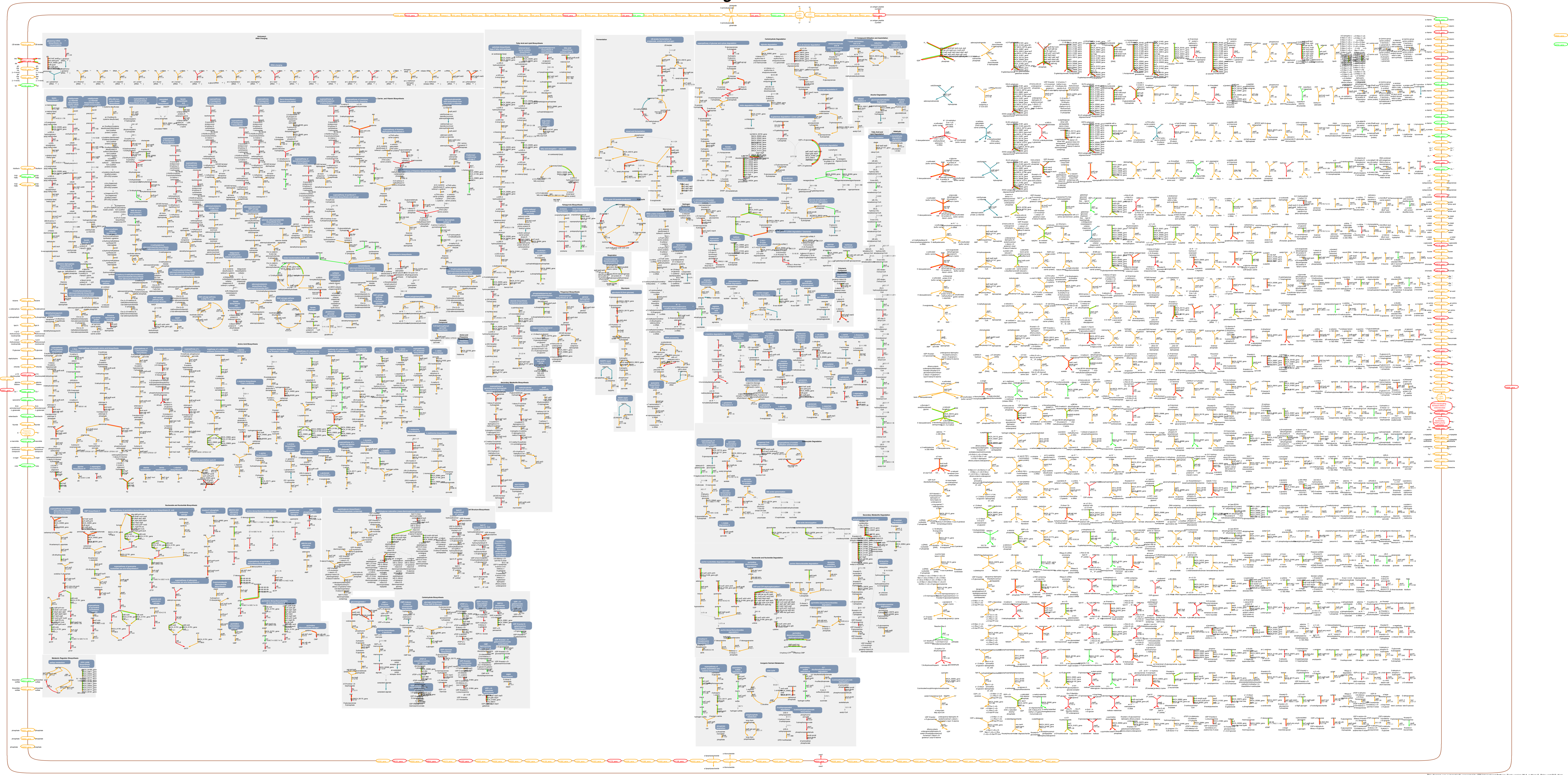

This diagram was automatically generated by SRI International Pathway Tools version 25.5, authors S. Paley and P.D. Karp.

### *Bacteroides thetaiotaomicron* Cellular Overview D5 vs. D2

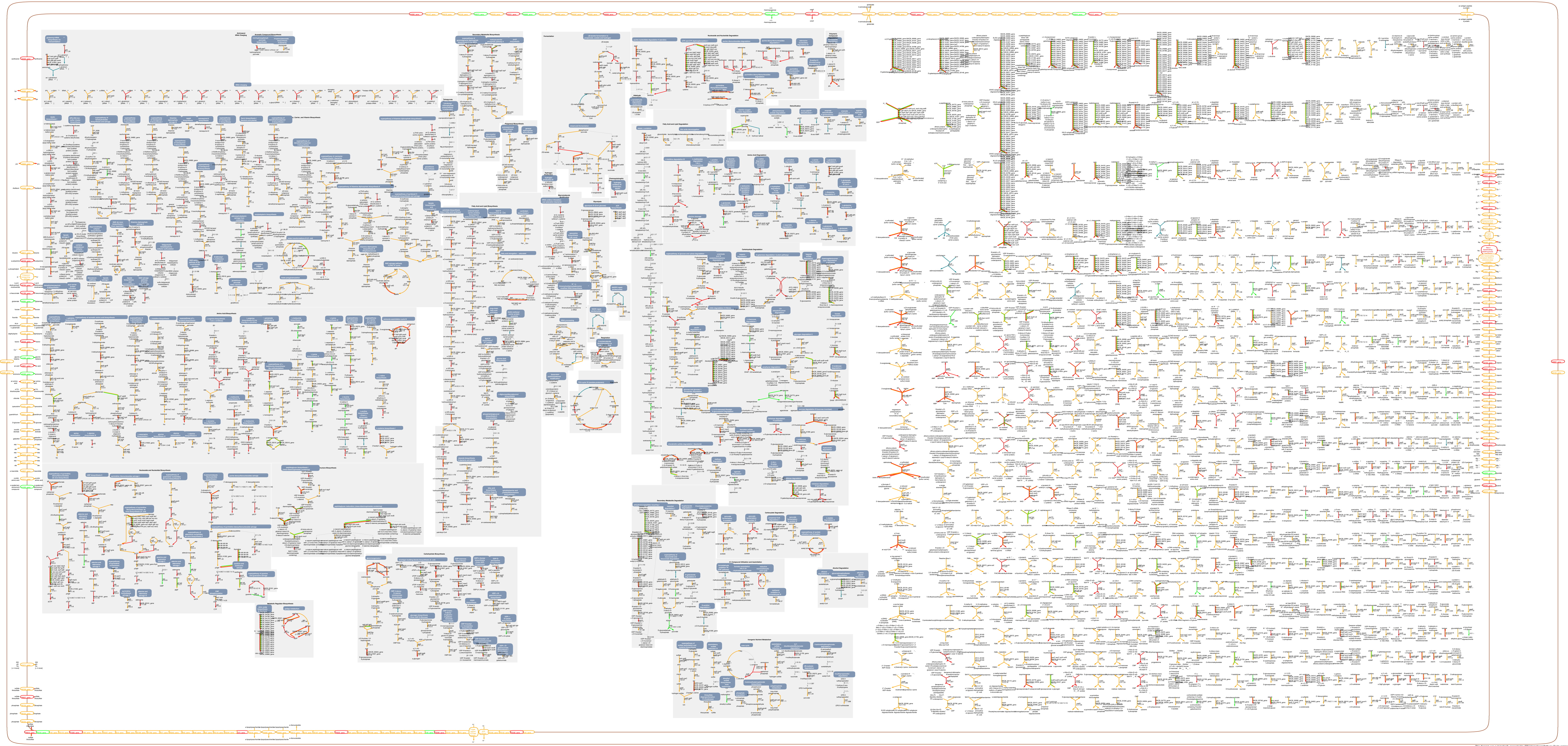


This diagram was automatically generated by SRI International Pathway Tools version 25.5, authors S. Paley and P.D. Karp.

### *Bacteroides* sp. Cellular Overview D5 vs. D2

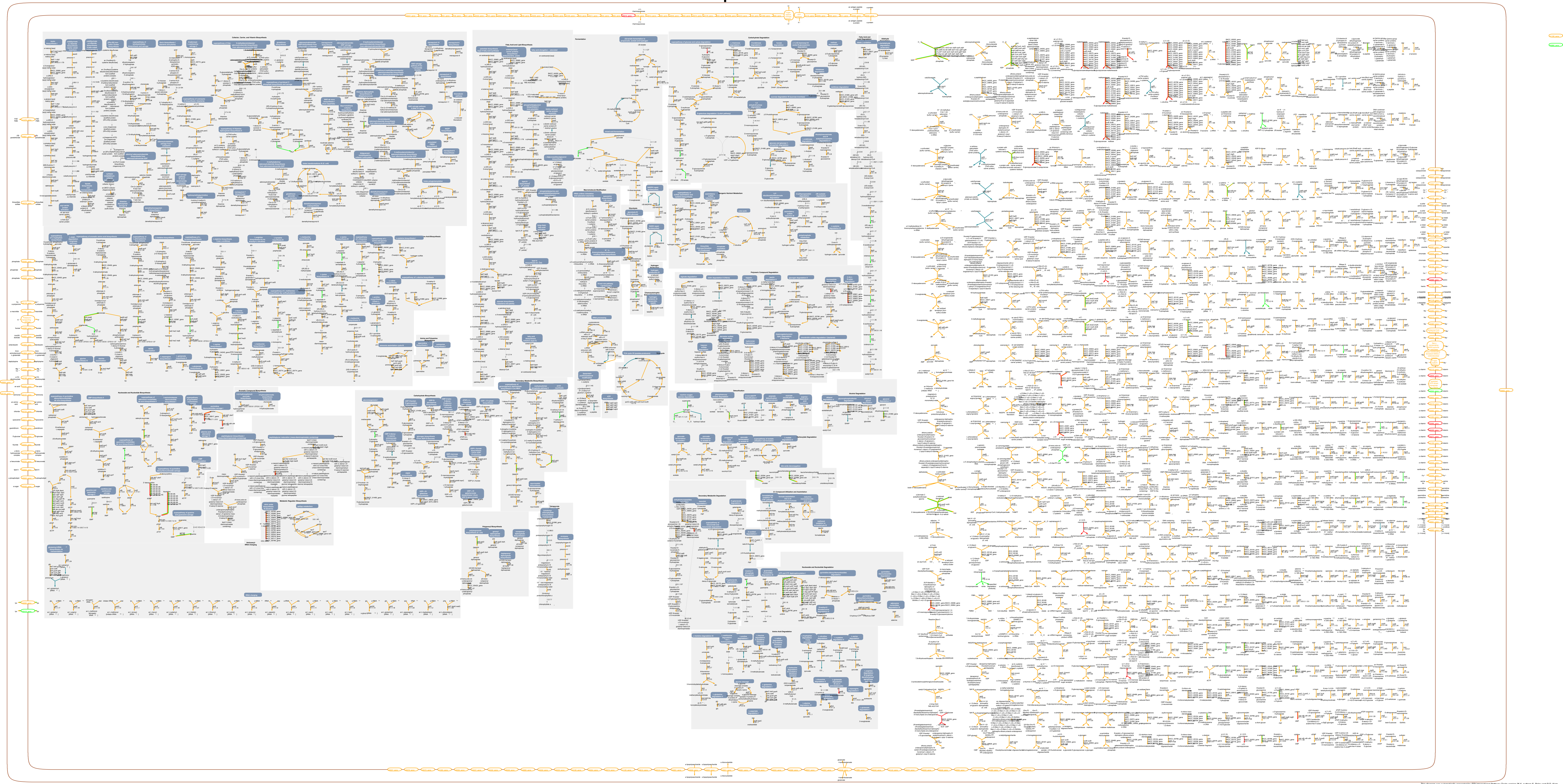

This diagram was automatically generated by SRI International Pathway Tools version 25.5, authors S. Paley and P.D. Karp.

### *Parabacteroides* sp. Cellular Overview D5 vs. D2

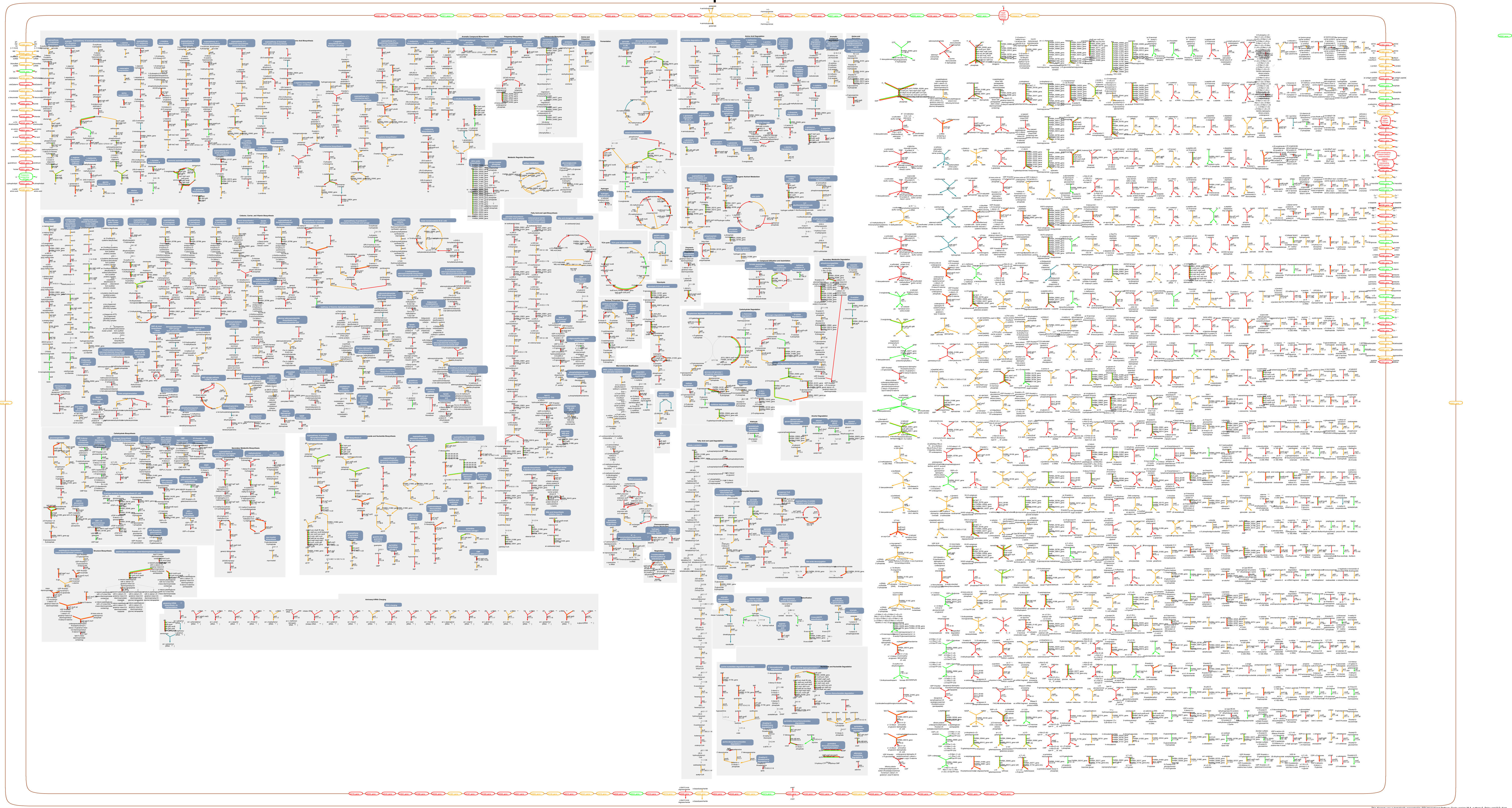

This diagram was automatically generated by SRI International Pathway Tools version 25.5, authors S. Paley and P.D. Karp.

### *Kerstersia gyiorum* Cellular Overview D5 vs. D2

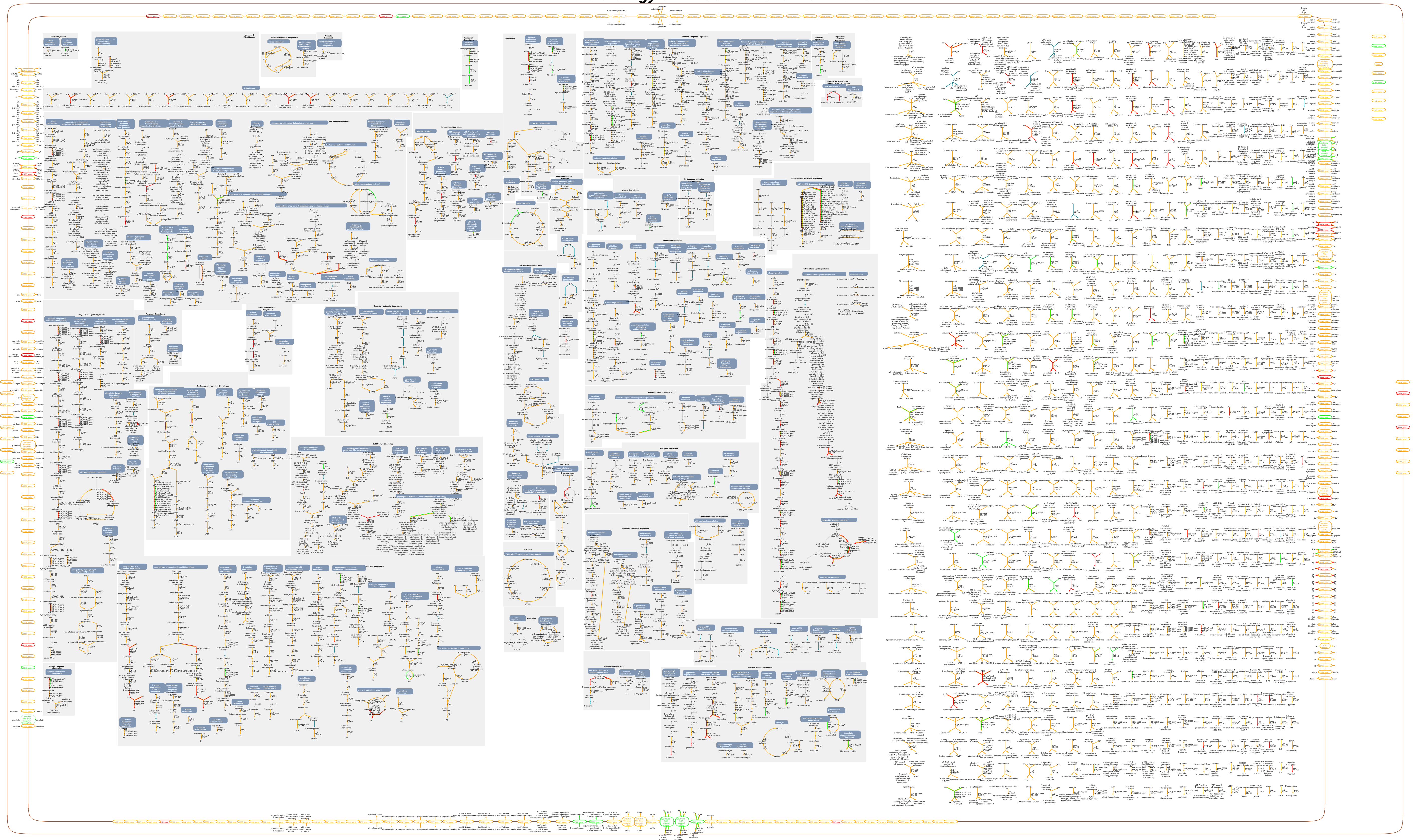

This diagram was automatically generated by SRI International Pathway Tools version 25.5, authors S. Paley and P.D. Karp.
